# Regulation of Nucleus Pulposus Cell Phenotype Through RhoA Signaling and Microenvironment

**DOI:** 10.64898/2026.04.05.716233

**Authors:** Gabriella Bond, Min Kyu M. Kim, Lauren Lisiewski, Timothy Jacobsen, Nadeen Chahine

## Abstract

Intervertebral disc degeneration is associated with loss of nucleus pulposus (NP) cell phenotype and extracellular matrix, both processes linked to changes in cytoskeletal contractility and cell shape. Here, we tested whether microenvironment-specific modulation of RhoA signaling can restore NP-like morphology and gene expression in NP cells cultured in 2D and in 3D alginate. In 2D monolayer culture, where cells are spread and mechanically activated, pharmacologic inhibition of RhoA with CT04 reduced RhoA activity, decreased actomyosin contractility gene expression, and shifted morphology toward a smaller, more circular phenotype. Bulk RNA sequencing showed that CT04 treatment increased expression of NP phenotypic and matrix-related genes including *ACAN*, *GDF5*, *CHST3*, and *MUSTN1* while decreasing expression of catabolic and fibroblast-associated genes including *ADAMTS1/9* and *COL1*, consistent with enrichment of extracellular matrix pathways. In contrast, RhoA activation with CN03 in 2D culture increased actin and phosphorylated myosin light chain intensity but produced limited phenotypic improvement. In 3D alginate, which minimizes integrin-mediated adhesion, baseline actomyosin markers were reduced relative to 2D culture. In alginate, RhoA activation with CN03 increased the amount of actin, phosphorylated myosin light chain, and actomyosin gene expression, yet also promoted a more compact, circular morphology and increased NP markers, including *ACAN* and *KRT19* with repeated dosing. Across culture conditions, increased cell roundness was consistently associated with increased *ACAN* expression, indicating strong coupling between cytoskeletal state, morphology, and NP matrix programs. Together, these findings demonstrate that RhoA pathway perturbation can promote NP phenotypic gene expression in both 2D and 3D culture, but the direction of optimal modulation depends on the microenvironment, supporting RhoA signaling as a context-dependent therapeutic target for disc regeneration.

## 1. INTRODUCTION

Low back pain (LBP) is the leading cause of disability worldwide, and intervertebral disc degeneration (IDD) contributes to up to 40% of LBP cases^1,2^. The intervertebral disc (IVD) is load-bearing tissue composed of the nucleus pulposus (NP), annulus fibrosus (AF), and cartilage end plates (CEP)^3^. During IDD, the IVD undergoes degradation of crucial extracellular matrix (ECM), structural disruption, and pathological load distributions^3–7^. Early detrimental effects of degeneration are typically observed in the NP, which is a hydrated and proteoglycan-rich tissue^4,8^. These microenvironmental changes are transmitted intracellularly through the mechanoresponsive NP cells^9,10^. This degenerative environment causes NP cells to lose their rounded morphology and develop cytoskeletal protrusions and diminishes healthy NP phenotype and anabolic ECM production^11,12^. Although major mechanotransduction pathways in NP cells have been identified, the context in which they control cell morphology and related cell phenotype are understudied.

Mechanotransduction pathways integrate external cues, such as matrix changes, into internal responses including cytoskeleton remodeling and downstream transcriptional changes^13^. The Rho/ROCK signaling pathway is a major mechanotransduction axis which regulates actomyosin contractility. In non-muscle cells, such as NP cells, RhoA signals molecule Rho-associated kinase (ROCK), which signals downstream targets that phosphorylate non-muscle myosin II^13,14^ (Figure 1). Non-muscle myosin II molecules are made up of three pairs of peptides: two heavy chains, two regulatory light chains that regulate the NMII activity, and two essential light chains which stabilize the heavy chain^15^. NMII has three isoforms, myosin IIA, IIB, and IIC, which are determined by the heavy chain, encoded for by *Myh9*, *Myh10*, and *Myh14*, respectively^15,16^. RhoA also promotes the polymerization of filamentous actin (F-actin), through diaphanous proteins and the inhibition of cofilin-mediated actin severing (Figure 1). The globular heads on NMII bind to actin, and form a contractile complex, which affects the cell cytoskeleton and, therefore, its morphology^13,14,17^. Downstream of this pathway are mechanosensitive transcription factors MRTF-A/SRF and YAP/TAZ. In NP cells, it has been shown that a rounded cell morphology keeps these coactivators in the cytosol^18^. When NP cells spread on a 2D surface, these factors are upregulated and localized to the nucleus^18^. However, deletion of either of these factors independently does not affect cell morphology. Return of NP rounded cell morphology was most prominent when changes occurred higher up in the signaling pathway, affecting actomyosin contractility, through substrate stiffness and F-actin depolymerization^18^. While the effects of actomyosin contractility on NP cell morphology has been studied, an understanding of their downstream effects on cell phenotype has not been established.

**Figure 1.**
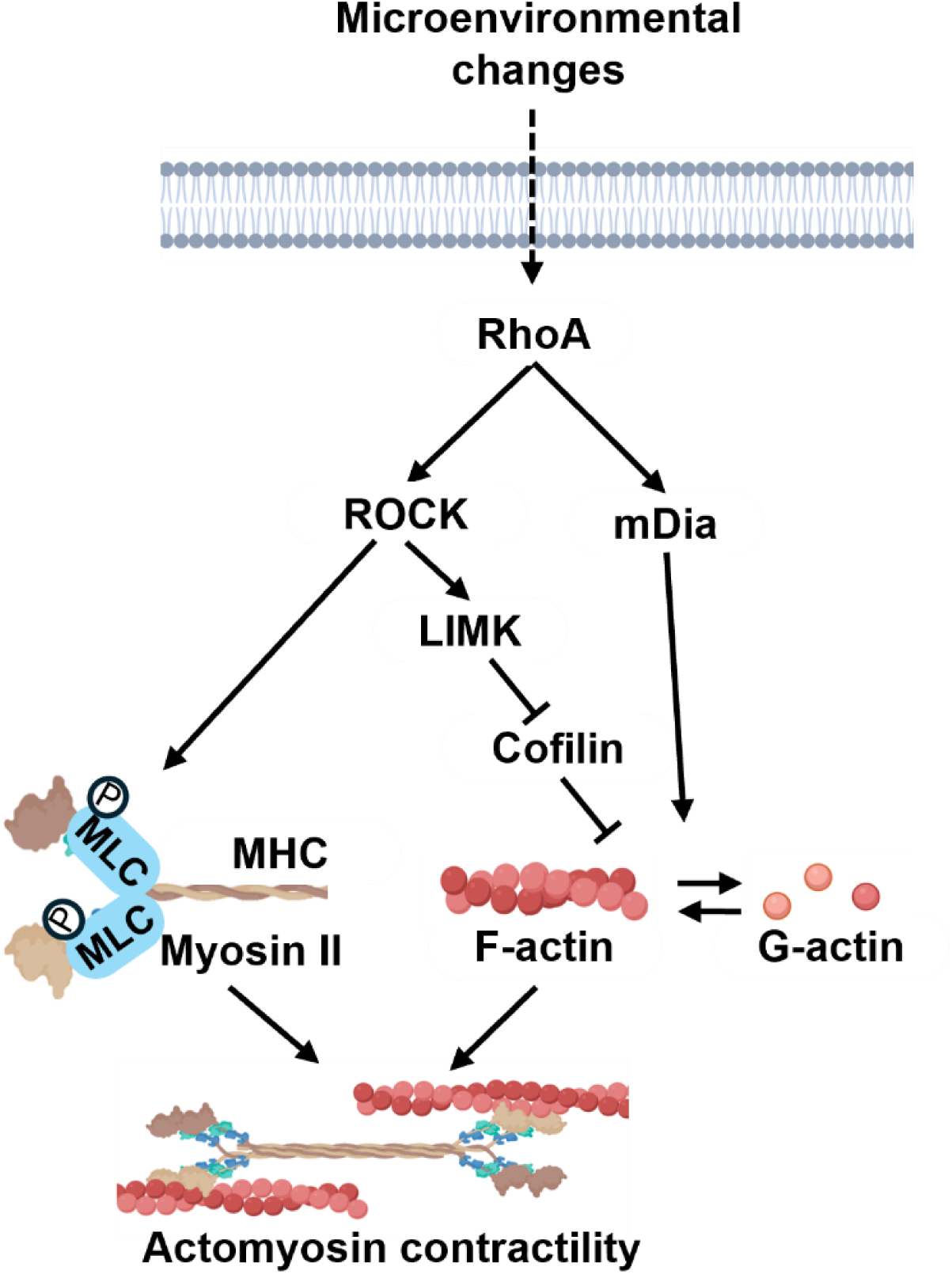
RhoA signaling pathway. RhoA signals downstream ROCK and mDia. ROCK increases phosphorylation of myosin light chain (MLC) and signals LIMK, which inhibits cofilin and allows for actin polymerization. mDia also influences actin polymerization. Myosin II and actin interact to alter cellular contractility.

Cell phenotype is highly dependent on cell morphology. Healthy NP cells are rounded, with a rich cortical actin cytoskeleton^19,20^. In contrast, NP cells cultured on tissue culture plastic form stress fibers, decrease GAG production, and progressively dedifferentiate with passaging^18,21,22^. Pharmacological disruption of F-actin reduces stress fibers, which is accompanied by an increase NP expression of aggrecan (*ACAN*) and collagen II (*COL2*)^18^. In other cell types, direct modulation of RhoA signaling alters both morphology and differentiation. In chondrocytes, inhibition of RhoA in 2D culture decreases cell area, which increases expression of chondrogenic genes *Gdf5* and *Sox6* and ECM genes *Acan*, *Col2,* and collagen VI (*Col6*) ^23^. Similarly, mesenchymal stromal cell (MSC) lineage commitment is influenced by shape, with rounded cells favoring chondrogenic/adipogenic fates and spread cells favoring osteogenesis^24–26^. NP cells are most studied in 3D hydrogel culture platforms since they better mimic *in situ* tissue environments. In soft hydrogel systems such as alginate, NP cells have decreased stress fibers compared to 2D culture and retain their natural morphology and phenotype^27,28^. Alginate is the most used hydrogel for culture of NP cells, due to its hydrated nature and tunable mechanical properties, supporting re-differentiation to their *in vivo* phenotype^29^. Additionally, it is a bioinert material, and while mechanotransduction is integrin-dependent in the NP in 2D and 3D culture, the lack of integrin binding sites reduces focal adhesions which promotes rounded morphology and NP phenotype^10,30,31^. However, this also creates a distinct mechanotransduction context compared with adhesive 2D culture. This phenotypic shift based on cell morphology is partially governed through the Rho/ROCK signaling pathway^25,32^. Rho/ROCK signaling effects on cell phenotypes are well-studied in muscle cells, such as fibroblasts and endothelial cells, but this relationship in nucleus pulposus cells has not been elucidated.

To evaluate the relationship between RhoA signaling and cell phenotype in different microenvironments, we cultured bovine NP cells in 2D monolayer or 3D alginate with RhoA perturbations. We activated RhoA signaling with CN03, and inhibited RhoA with CT04, then assessed cell morphology and associated changes in gene expression. We find that RhoA inhibition in 2D culture and RhoA activation in 3D culture both decrease cell area and increase expression of NP genes, highlighting microenvironment-dependent regulation of NP morphology and phenotype by RhoA signaling.

## 2. MATERIALS AND METHODS

### 2.1 Materials

Phosphate buffered saline (PBS), Penicillin-Streptomycin (PS), high glucose Dulbecco’s modified Eagle’s medium (DMEM), 16% formaldehyde, and Tween-20, were purchased from Thermo Fisher Scientific (Waltham, MA). Premium select fetal bovine serum (FBS) was purchased from R&D systems. Collagenase type I (LS004194) and type II (LS004174) were purchased from Worthington Biochemical. Dimethyl sulfoxide (DMSO), alginic acid sodium salt, sodium chloride (NaCl), HEPES, calcium chloride (CaCl_2_), bovine serum albumin (BSA), magnesium sulfate (MgSO_4_) and triton-X100 were purchased from Sigma-Aldrich. Hank’s Balanced Salt Solution (HBSS) was purchased from Corning. Rho Activator II (CN03), Rho Inhibitor I (CT04), G-LISA 6 RhoA Activation Assay Biochem Kit (BK124), and Total RhoA ELISA (BK150) were sourced from Cytoskeleton, Inc. (Denver, CO). RNeasy Mini and Micro kits for RNA isolation was sourced from Qiagen. RT-qPCR primer sequences were purchased from Integrated DNA Technologies. Rabbit-αPMLC (3674s) antibody was sourced from Cell Signaling, Goat αRabbit-Alexa 488 (ab150081) antibody was sourced form Abcam, Zombie Red Cell Viability Dye (423109) was sourced from Biolegend, and Alexa Fluor 647 Phalloidin (A22287) and DAPI (62248) nuclear stain were both sourced from Thermo Fisher Scientific. LIVE/DEAD Cell Viability Assays purchased from Thermo Fisher Scientific.

### 2.2. Bovine nucleus pulposus (bNP) cell isolation

Lumbar spines (n=5) from juvenile (3-week-old) bovine animals (male) were shipped overnight on ice from an abattoir (Research 87). Upon arrival, the spine was cleaned of extra tissue and sterilized in iodine solution for an hour before use in the hood. IVDs were dissected from the spinal column and NP and AF were separated and stored in PBS + 10% PS. The tissue was pooled and digested in complete media [high glucose DMEM + 10% FBS + 1% PS] supplemented with collagenase type I (0.3 mg/ml) and collagenase type II (0.3 mg/ml) for 3 hours at 37°C with gentle agitation. bNP cell digest was passed through a 70-μm cell strainer, washed, counted, and plated to be expanded in in complete medium. Cells were frozen at P1 in 10% DMSO in complete media and stored in liquid nitrogen until use.

### 2.3. 2D Cell Culture

Cells were thawed from frozen and expanded at P1, then lifted and replated at P2 in 12-well plates at a density of 200,000 cells per well. The cells were cultured for 24 hours in complete media. Cells were treated with the following groups: (i) CN03, RhoA activator at 0.2 μg/ml, 1 μg/ml, 5 μg/ml and (ii) CT04, RhoA inhibitor at 0.5 μg/ml and 2 μg/ml.

### 2.4. Alginate Bead Synthesis

A 1.2% alginate solution was created by dissolving alginic acid sodium salt in 150mM NaCl + 10mM HEPES overnight. Cells were suspended at a concentration of 1 million cells per milliliter of alginate. Beads were created by passing the mixture through a 21G needle dropwise into a bath of 102 mM CaCl_2_ and allowed to crosslink for 15 minutes at 37°C. The cells in beads were cultured for 24 hours in complete media before treatment. Cells in 3D were treated for 24 hours (1 day) with CN03 at 1 μg/ml. In 3D, cells were also treated with CN03 (1 μg/ml) over a 7-day period using either: (i) a single dose, where CN03 was added on day 1 and removed upon media change at day 3, or (ii) daily dose, where CN03 was added daily on days 1 through 6. Cells in alginate beads were collected for assays at day 7.

### 2.5. RhoA Activity Quantification

Cells taken from 2D culture were placed on ice and washed with ice-cold PBS one time. They were lysed using scraping with lysis buffer supplemented with protease inhibitor provided in the ELISA kits. Lysates were collected and clarified by centrifugation at 10,000g at 4°C for 1 minute. Total protein levels were quantified using Precision Red^TM^ Advanced Protein Assay Reagent, and the rest of the samples were snap-frozen. Prior to use in the ELISA, lysates were equalized to the same protein concentration within the range of 0.4-0.8 mg/ml. Total RhoA levels were measured using the Total RhoA ELISA kit and GTP-bound RhoA levels were measured using the RhoA G-LISA Activation Assay kit according to the manufacturer’s protocol.

### 2.6. RNA Isolation and Gene Expression

After the designated culture period, cells in 2D were lysed and RNA was isolated using the RNeasy Mini Kit according to the manufacturer’s protocol. To collect RNA from cells in 3D, alginate beads were digested in sodium citrate (50mM sodium citrate, 140mM NaCl, 10 mM HEPES in water) for 15 minutes with agitation. Cell pellets were collected after spinning digests down at 500g for 5 minutes and then washed with PBS and pelleted again. Cell pellets were resuspended in TRIZol and RNA was isolated using the RNeasy Micro kit according to the manufacturer’s protocol. RT-qPCR was conducted to measure levels of gene expression in actomyosin contractility genes (*MYL6, MYL9, MYH9, MYH10, ACTA2*), ECM genes (*ACAN, COL1)*, and NP phenotypic marker (*KRT19).* Primer sequences can be found in table 1.

**Table 1.**
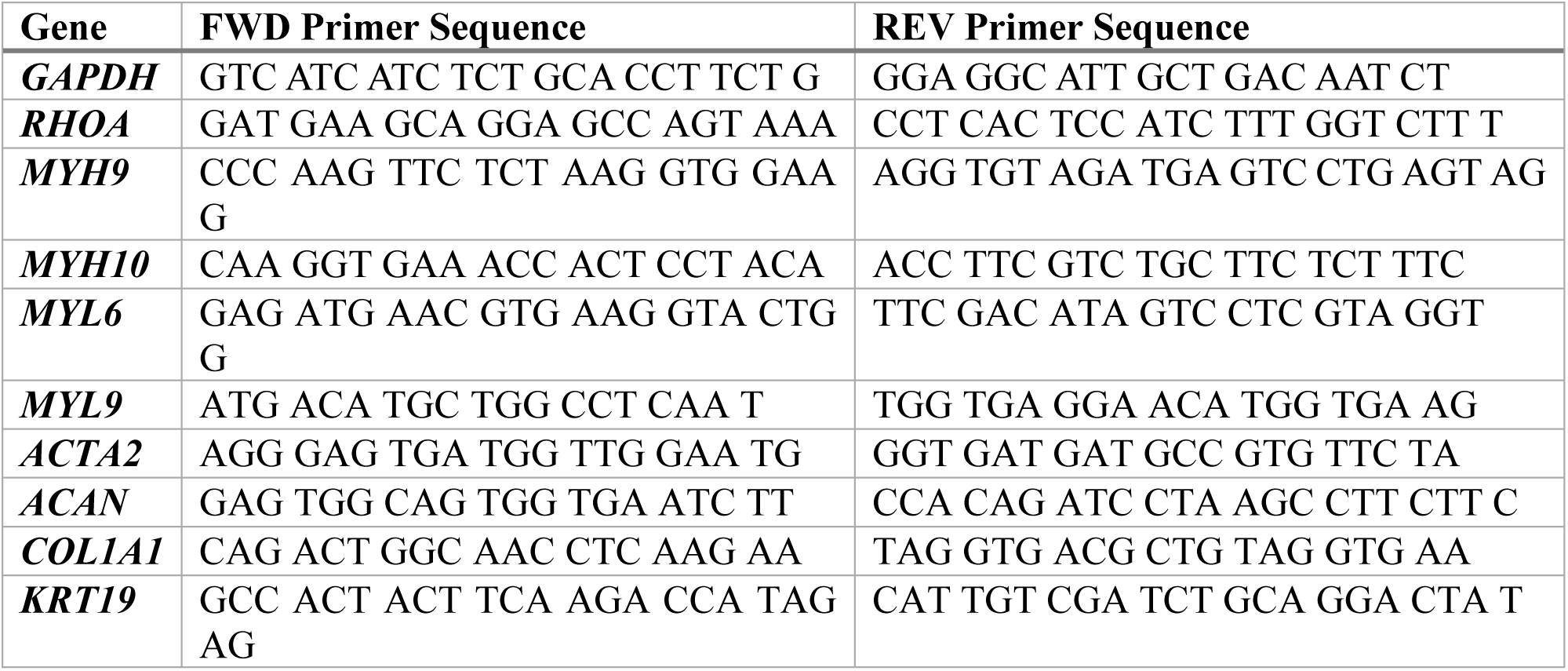
Primer sequences for RT-qPCR.

### 2.7. Bulk Transcriptomics

Isolated RNA from cells in 2D were used for bulk RNA sequencing conducted by the Columbia Genome Center using an AVITI Instrument. For each sample, library preparation was completed with STRPOLYA kits and sequenced on Illumina NovaSeq 6000. In R, the variance stabilizing function in DESeq2 was used to normalize gene count data. Differentially expressed genes (DEGs) were identified based on |log2FoldChange| ≥ 1.5 and adjusted p value (padj) < 0.05. To create PCA plots, variance stabilizing functions were performed. Over-representation analysis was performed for each treatment group vs. control using web-based EnrichR, which identified major signaling pathways of these DEGs. Leading-edge genes were also extracted pathways of interest in both CT04 and CN03 treated groups. To uncover subtle changes in signaling, Gene Set Enrichment Analysis (GSEA) using the Reactome gene set was performed in R, with p<0.05 cutoff for pathway significance. Subsets of different pathways were displayed in ranked form using Normalized Enrichment Score (NES).

### 2.8. Immunostaining and Imaging

After treatment in 2D culture, cells were rinsed with PBS and fixed in 4% paraformaldehyde (PFA) at room temperature for 30 minutes. Cells were rinsed with 0.05% Tween-20 in PBS (PBS-T) three times and then permeabilized with 0.5% Triton-X100 in PBS for 10 minutes at room temperature. Cells were rinsed again with PBS-T three times and blocked with 1% BSA in PBS-T for an hour at room temperature. Primary antibody for phosphorylated myosin light chain (pMLC) (1:100 in 1% BSA) was added to cells and left overnight at 4°C. The primary antibody was removed, cells were rinsed with PBS-T 3 times, and the secondary antibody for pMLC was added (1:500 in PBS-T) along with the conjugated stain for actin (1:1000 in PBS-T), for an hour at room temperature protected from light. Cells were rinsed 3 times with PBS-T and then DAPI (1:1000 in PBS-T) was added for 30 minutes. Samples were rinsed twice with PBS-T and imaged under an inverted lens confocal. Images were analyzed using a custom pipeline built in CellProfiler to quantify actin mean intensity, pMLC mean intensity and cell area.

After treatment in 3D culture, cells were collected and washed with Hank’s Balanced Salt Solution (HBSS) supplemented with 1.26mM of CaCl2 and 400uM of MgSO4. All wash steps were 5 minutes at room temperature with HBSS + Ca/Mg to ensure full penetration of the alginate bead. Cells were then stained with Zombie Red Dye (1:2000 in HBSS) for cell viability for 30 minutes at RT. All subsequent steps occurred with minimal light exposure. Cells were washed with HBSS + Ca/Mg and fixed in 4% PFA for 1 hour at room temperature. Cells were rinsed again twice and permeabilized with 0.5% Triton-X for 15 minutes at room temperature. Cells were rinsed again and then blocked with 1% BSA for 1 hour at room temperature. Staining for actin and pMLC in 3D followed the same procedure described for NP cells in 2D. After these steps, the cells were removed from the alginate beads using a sodium citrate (50mM sodium citrate, 140mM NaCl, 10 mM HEPES in water) dissolution for 15 minutes on a shaker at room temperature. Cells were spun down at 600g for 5 minutes, washed with PBS, and then spun down again at the same speed. Cells were resuspended in at least 35μl of PBS with DAPI (1:1000). Cells were images on the Cytek^®^ Amnis^®^ ImageStream^®X^ Mk II imaging flow cytometer and analyzed using the IDEAS^®^ software for mean intensity of actin and pMLC, as well as cell shape parameters (cellular area and circularity). Cells were gated for in-focus, live cells, and single nuclei.

### 2.9. Statistics

Quantitative data are reported as mean ± standard deviation. RhoA ELSIA was conducted with n = 4-5 technical replicates. Gene expression data is based on n = 2-3 biological replicates. RNA sequencing studies were conducted with n = 3 biological replicates. Comparisons among treatment groups that had parametric data were conducted with one-way ANOVA with Tukey post-hoc. Data with both treatment and time point were compared using two-way ANOVA with Tukey post-hoc. Comparisons among treatment groups with non-parametric data were conducted using Kruskal-Wallis with Dunn’s post-hoc. Singular parametric comparisons were conducted using Student’s t-test. Statistical analysis was performed in Graphpad Prism with significance defined as a p-value < 0.05.

## 3. RESULTS

### 3.1 CN03 activates RhoA and CT04 inhibits RhoA in NP cells culture in 2D in a dose dependent manner

CN03 treatment of cells in 2D culture increased RhoA activity in a dose-dependent manner, where normalized active RhoA content was significantly increased with 1μg/ml and 5μg/ml compared to untreated. Treatment with 5μg/ml CN03 increased gene expression levels of *MYH9, MYL6, MYL9,* and *ACTA2,* while 1μg/ml CN03 increased expression of *MYL6, MYL9,* and *ACTA2* compared to untreated (Figure 2A). Cells dosed with different concentrations of CT04 only exhibited a significant decrease in active RhoA content at the higher dose, 2μg/ml. However, this data was not normalized to total RhoA content for clarity, since both CT04-treated groups also had a significant decrease in total RhoA (Figure 2B). To confirm that the cell viability was unaffected, we checked total protein content and performed live-dead staining on the cells. Results show that there were no differences due to CT04 treatment (Figure S1). Both doses of CT04 (0.5μg/ml, 2μg/ml) decreased expression of *MYH9, MYH10, MYL9,* and *ACTA2,* while only the higher dose of CT04 decreased expression of *MYL6* (Figure 2B). From these preliminary results, 1μg/ml of CN03 and 2μg/ml of CT04 were chosen for subsequent studies.

**Figure 2.**
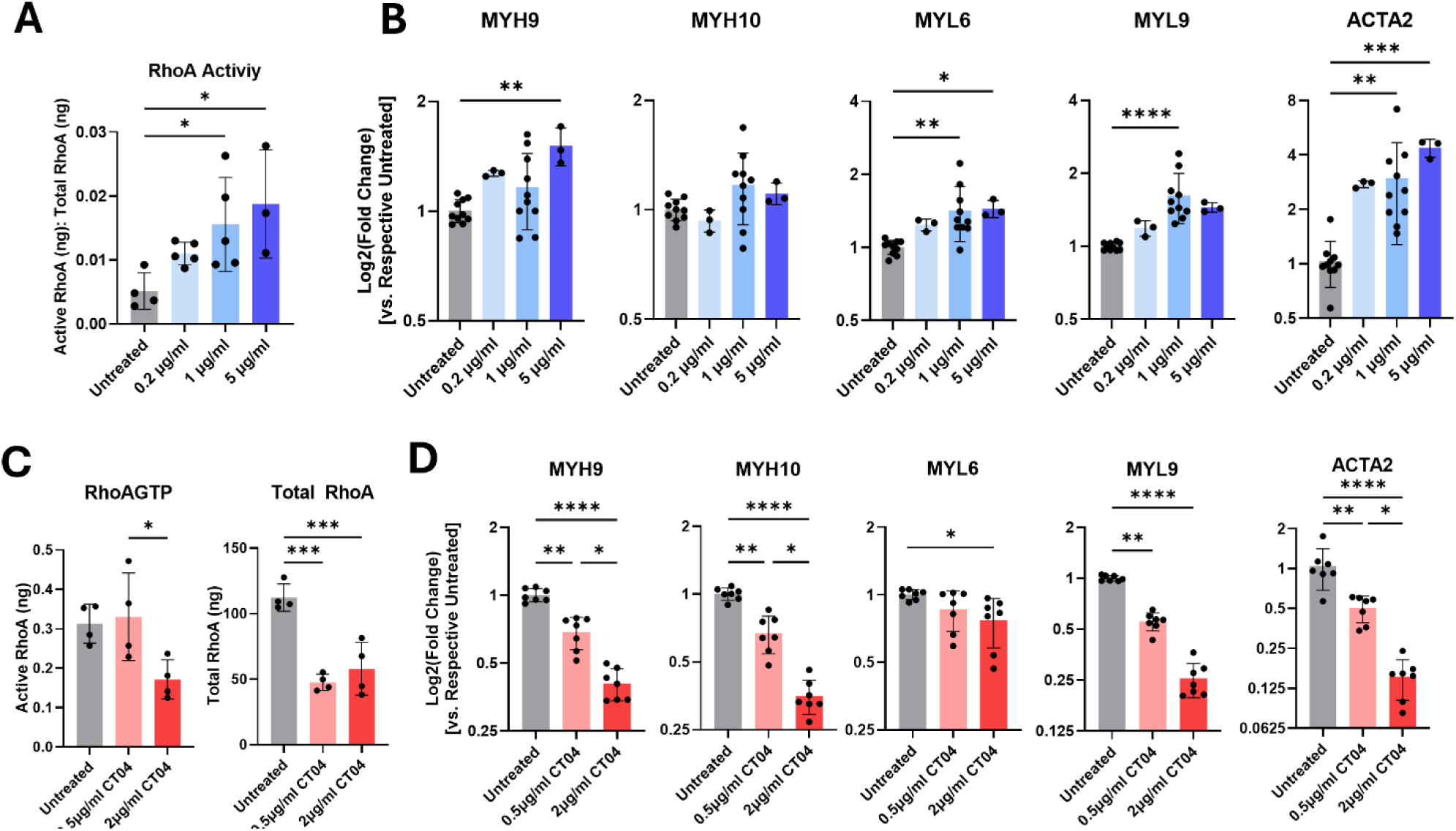
Dose response of NP cells in 2D to CN03 and CT04 treatment. Higher doses of CN03 A) increased RhoA activity and B) increased expression of actomyosin genes in NP cells in 2D. CT04 treatment C) decreased active (2μg/ml only) and total RhoA content and D) decreased expression of actomyosin genes. *p<0.05, **p<0.01, ***p < 0.001, **** p<0.0001

### 3.2 In 2D, RhoA activation and inhibition both affect cytoskeleton, but only RhoA inhibition affects morphology

In 2D culture, CN03 treatment increased mean intensity of actin and pMLC in NP cells, however, it did not significantly change cell area compared to untreated (Figure 3). In CT04-treated groups, NP cells showed lower pMLC intensity and actin intensity. Untreated cells cultured in 2D exhibit a spread morphology, but they become more circular when treated with 2 μg/ml of CT04, as evidenced by the lower cell area (Figure 3).

**Figure 3.**
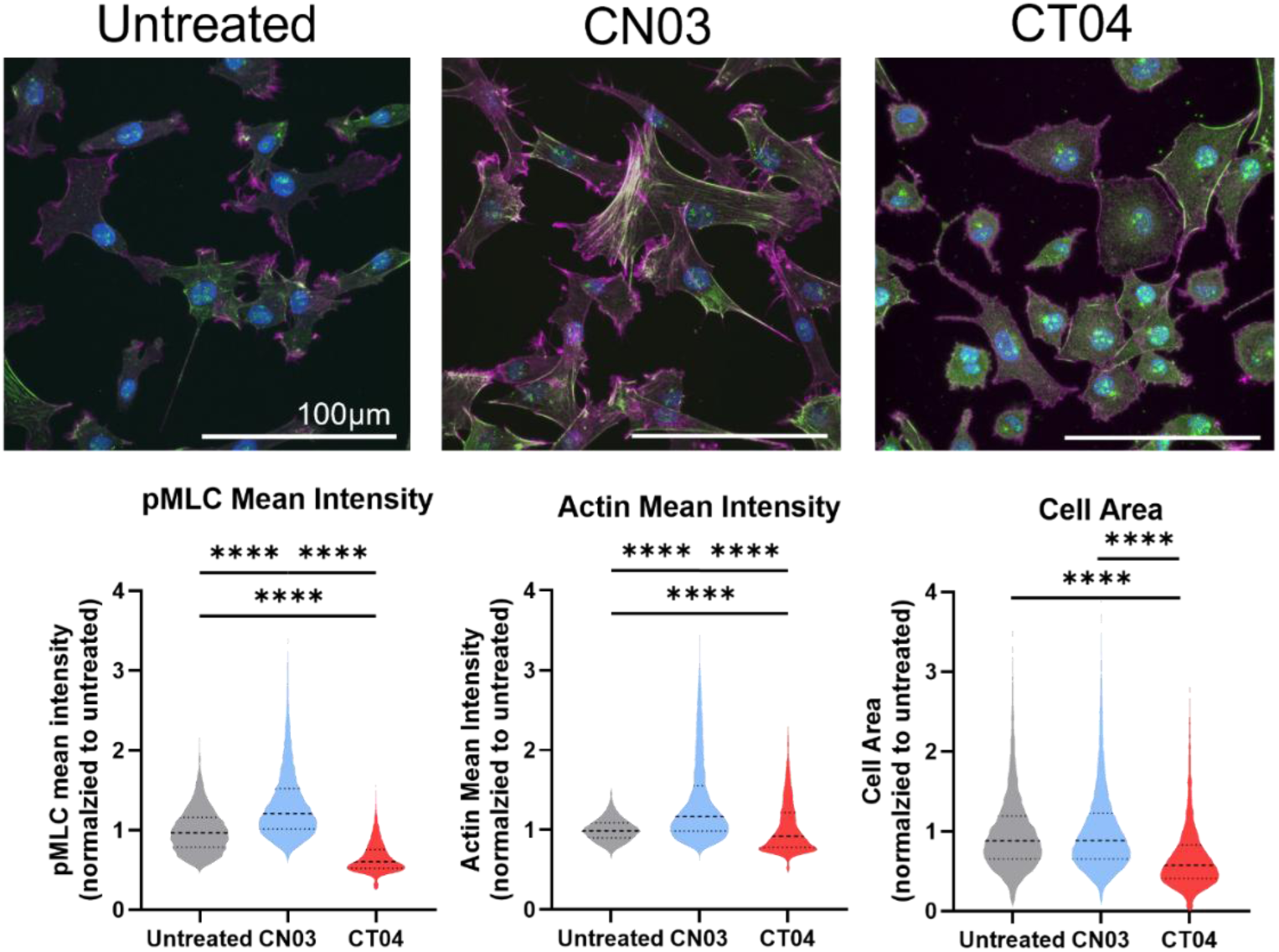
2D Imaging of cells treated with CN03 and CT04. Treatment with CN03 increased pMLC and actin intensity but did not affect cell area. Treatment with CT04 decreased pMLC intensity, actin intensity, and cell area. ****p<0.0001

### 3.3 RhoA activation in 2D dysregulates ECM and integrin signaling

A principal component analysis was applied to the differentially expressed genes (DEGs) based on RNA sequencing. CN03-treated groups clustered in the PCA plot, with 191 DEGs total (Figure 4A). Over-representation EnrichR analysis using these DEGs showed that the top 15 implicated pathways are related to cholesterol biosynthesis and steroids, G protein-coupled receptor (GPCR) signaling, Rho signaling, and ECM organization (Figure 4B). To elucidate what DEGs contribute the ECM organization signaling pathways, we plotted their fold change. The expression of ECM molecules *COL6A6* and *LAMA3*, integrins (*ITGA7* and 8), and degradative gene *MMP17* were upregulated with CN03 treatment (Figure 4B). In contrast, ECM molecules *COL27A1* and *COL21A1* were downregulated along with integrin *ITGA9*, degradative gene *ADAMTS3, a*nd growth factor *GDF5* (Figure 4B). When using GSEA reactome, which uncovers more subtle changes by using the entire gene list, Rho GTPase signaling pathways were positively enriched, along with integrin signaling (Figure 4C). The same GPCR signaling pathways observed in EnrichR analysis were negatively enriched in Reactome GSEA (Figure 4C).

**Figure 4.**
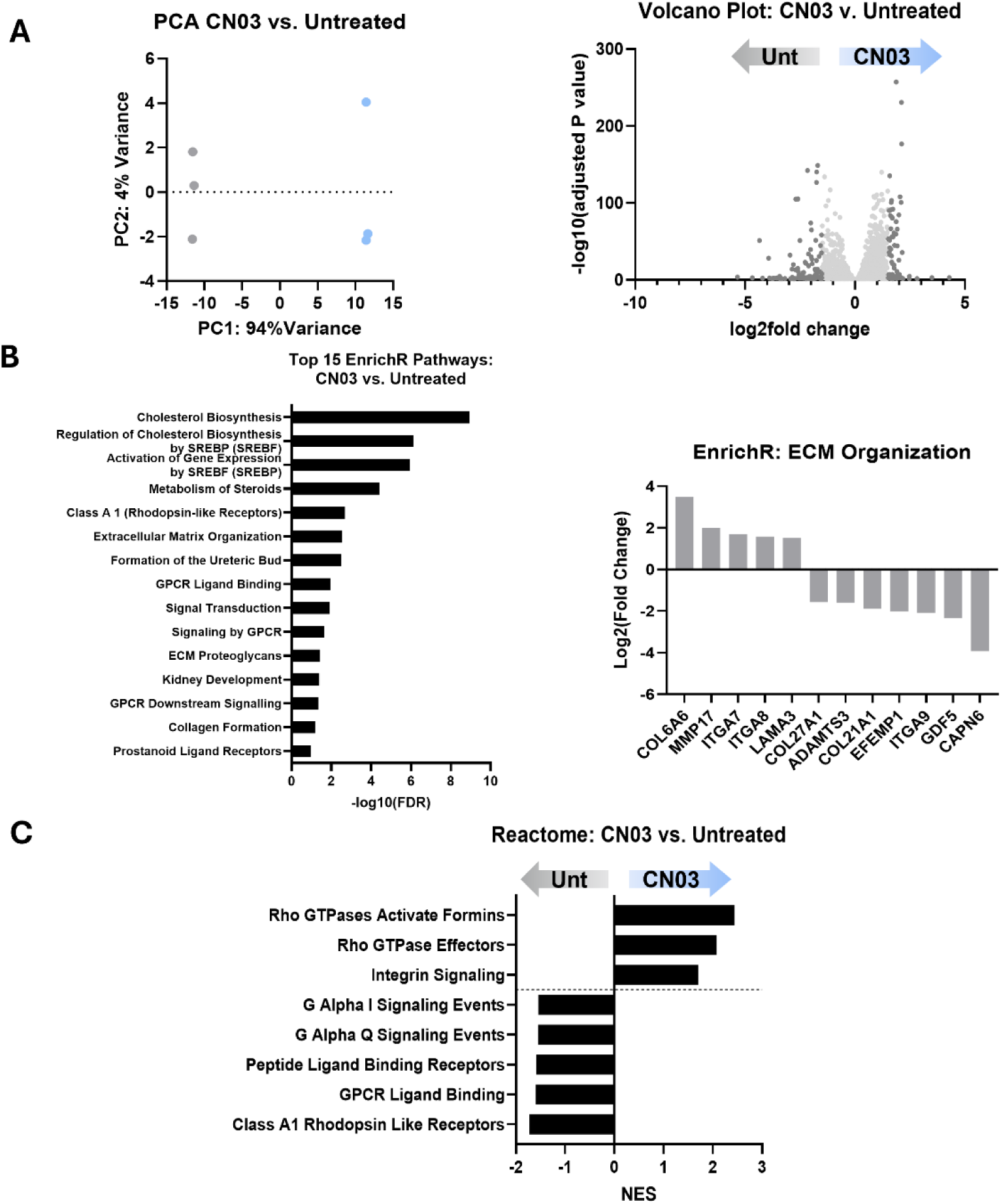
RNA sequencing of NP cells treated with CN03. A) CN03-treated NP cells clustered from the untreated cells in the PCA plot, with 191 DEGs in the volcano plot. B) Top 15 EnrichR pathways showed cholesterol synthesis, ECM organization, and GPCR signaling. The ECM organization pathway had 12 contributing DEGs. C) GSEA using Reactome showed upregulation of Rho GTPase pathways and downregulation of GPCR signaling pathways.

### 3.4 RhoA inhibition in 2D promotes NP ECM and phenotypic gene expression

The morphological change observed in NP cells in 2D treated with CT04 groups is accompanied by changes in bulk RNA sequencing data. First, there was distinct and separable clustering between the untreated and CT04 treated groups on the PCA plot (Figure 5A). Out of the 739 DEGs in CT04 vs. untreated NP cells, some of the top 25 genes ranked by adjusted p-value from DESeq2 were *GDF5, CHST3,* and *MUSTN1* (Figure 5A). Over-representation EnrichR analysis on these DEGs showed enrichment of signaling pathways within the broad categories of cell cycle, ECM organization, and signal transduction, including Rho signaling (Figure 5B). To understand the changes in the ECM pathway, we further examined the DEGs contributing to this pathway. Out of the 36 genes, there was upregulation of NP ECM components (*BCAN*, *ACAN*, and *LAMB3*), growth factors (*GDF5*, *TGFB2*, *BMP2*), integrin *ITGA9*, and degradative molecules (*MMP3*, *MMP15*, *MMP24*) (Figure 5B). There was also downregulation of integrin *ITGA8,* degradative molecules *ADAMTS1* and *ADAMTS9,* and cell-adhesion gene *ICAM1* (Figure 5B). Reactome GSEA produced 148 pathways, where we confirmed that the CT04 inhibited RhoA signaling pathway with significant negatively enriched pathways: Rho GTPase effectors, smooth muscle contraction, and Rho activates formins (Figure 5C). Additionally, there were many positively enriched pathways including collagen formation, ECM organization, and ECM proteoglycans (Figure 5C). To further examine genes that were relevant in NP ECM, we conducted PCR and found that there was an increase in *ACAN* expression accompanied by a decrease in *COL1* expression (Figure 5D).

**Figure 5.**
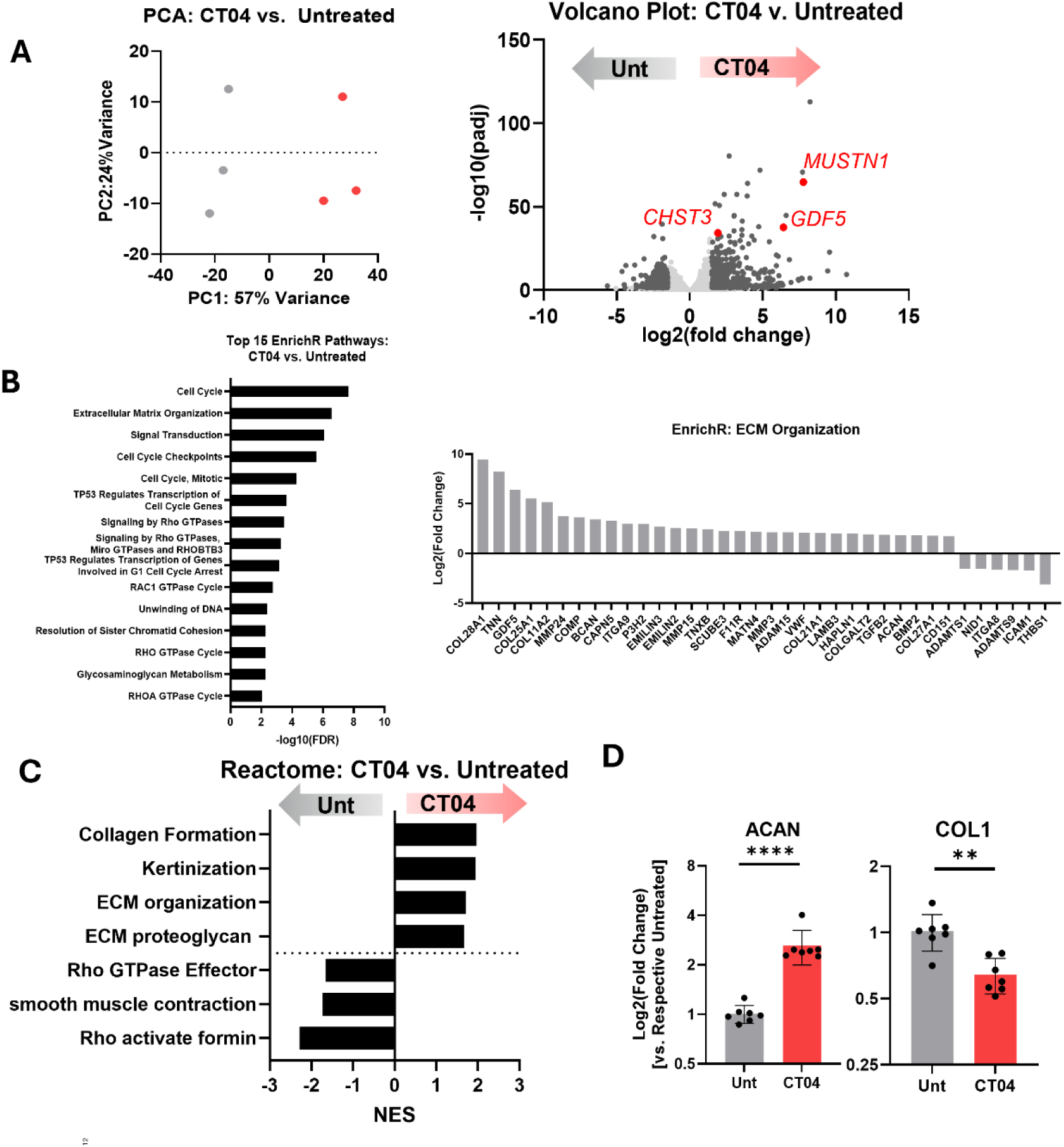
RNA sequencing of NP cells treated with CT04. A) PCA plot of CT04-treated NP cells vs. untreated cells, which clustered from each other with 739 DEGs in the volcano plot. B) Top 15 EnrichR pathways showed enrichment of pathways related to cell cycle, ECM organization, and Rho GTPases. The 36 DEGs contributing to ECM organization ranked by log2(fold change) are plotted. C) GSEA Reactome revealed upregulation of ECM and keratinization pathways and downregulation of Rho GTPase and contraction pathways. D) PCR analysis revealed a two-fold increase of *ACAN* expression and a decrease in *COL1* expression. **p<0.01, **** p<0.0001

**Figure 6.**
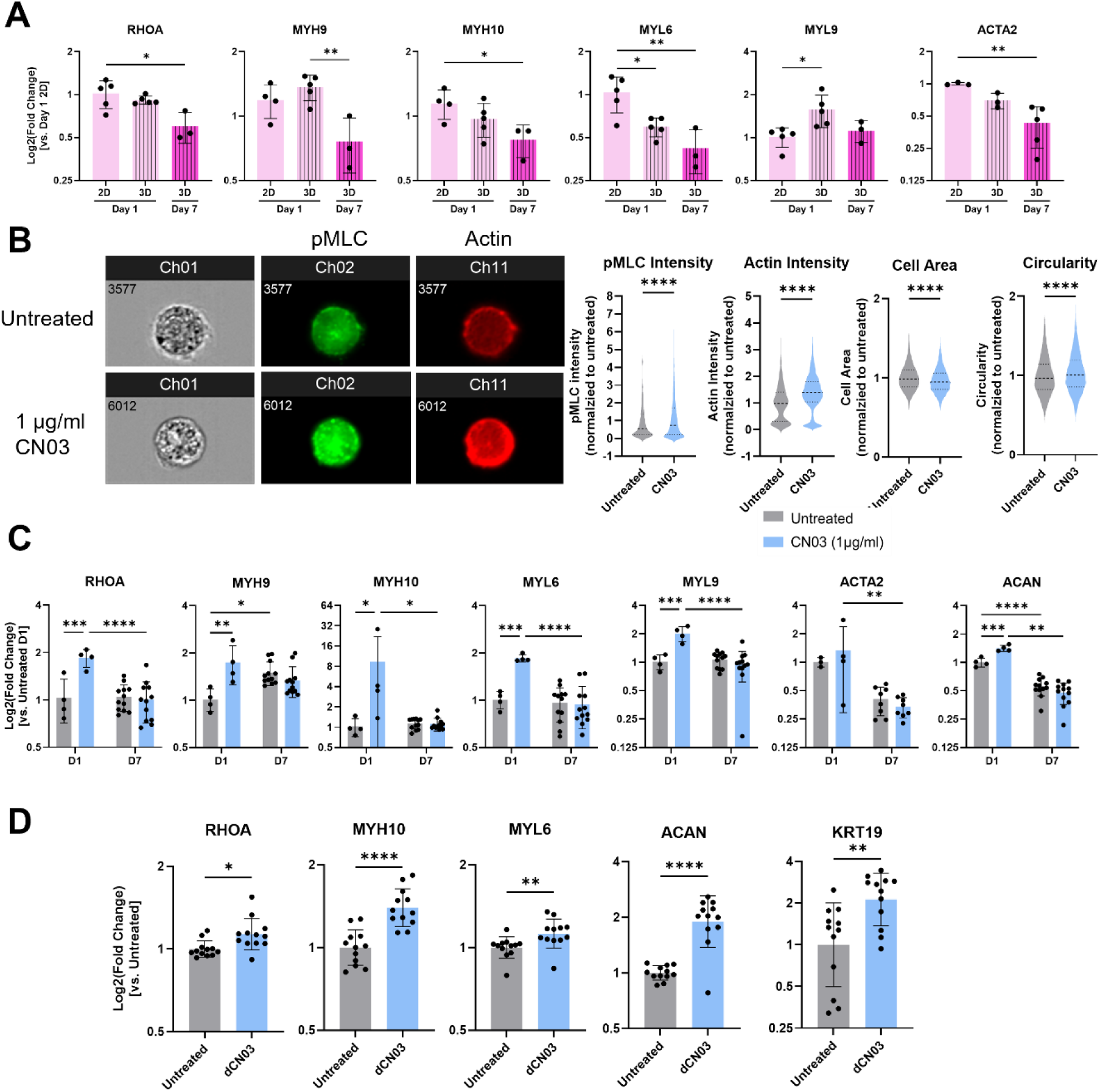
RhoA signaling and CN03 treatment of NP cells in alginate for 1- and 7- days. A) Over time in 3D alginate culture, expression of actomyosin genes decreased. B) Imaging flow cytometry analysis showed that 1-day CN03 treatment increased pMLC and actin intensity, decreased cell area, and increased cell circularity. C) 1-day CN03 treatment increased actomyosin and *ACAN* gene expression, but this was not sustained out to 7 days. D) Daily addition of CN03 for 7 days increased *RHOA*, *MYH10*, *MYL6*, *ACAN*, and *KRT19* expression compared to untreated. *p<0.05, **p<0.01, ***p<0.001, ****p<0.0001

### 3.5 CN03 treatment recovers loss of RhoA activity and promotes NP ECM in 3D alginate culture

Untreated NP cells cultured in 3D alginate beads for 1 day showed decreased expression of *MYL6* and *ACTA2* when compared to NP cells in 2D culture. At 7 days, NP cells in alginate culture exhibited decreased expression levels of *RHOA, MYL6, MYH9, MYH10,* and *ACTA2* when compared to cells in 2D after 1 day. Over time in alginate, from 1 day to 7 days, expression of *RHOA* and *MYH9* significantly decreased (Figure 5A). CN03-treatement of NP cells in alginate in 3D increased intensity levels of actin and pMLC (Figure 5B). This validated that the selected dose of CN03 used in 2D culture is effective in 3D culture systems. In addition, these cells had decreased cell area and increased cell circularity (Figure 5B). When compared to their untreated counterparts, CN03-treated NP cells in 3D exhibited increased gene expression of *RHOA, MYL6, MYL9, MYH9,* and *MYH10* and more notably, *ACAN* (Figure 5C). However, when this dose was removed after 1 day and cells are maintained in culture for 7 days total, this effect was diminished. There was no difference between the untreated and CN03-treated groups with a single dose at 7 days (Figure 5C), highlighting that RhoA activation is transient. Therefore, we decided to dose the cells in a sustained fashion, by administering a daily dose of CN03 into the cell culture media. NP cells that were treated daily with CN03 for 7 days exhibited increased gene expression of *RHOA, MYL6,* and *MYH10* compared to untreated (Figure 5D). In addition, there was an increase in NP-markers *ACAN* and *KRT19* (Figure 5D). This highlights that it is necessary for continuous delivery of CN03 to NP cells to sustain its benefits.

## 4. DISCUSSION

The goal of the study was to assess the effects of RhoA pathway on NP cell morphology and cell phenotype in 2D and 3D microenvironments. We find that promotion of NP cell phenotype can be achieved by perturbation of the RhoA pathway in a microenvironment-specific manner. In 2D, RhoA inhibition of flattened NP cells decreased RhoA activity, actomyosin gene expression, cell area and increased circularity, pushing NP cells towards a more physiologically relevant morphology. In 3D, when cells start out in rounded morphology, RhoA activation increased actomyosin gene expression, actin intensity, and pMLC intensity leading to decreased cell area and increased circularity. In both microenvironments, the promotion of rounded cell morphology was accompanied by increases in expression of pro-NP genes, such as *ACAN, GDF5, CHST3,* and *MUSTN1* in 2D, and *ACAN* and *KRT19* in 3D. These findings demonstrate that perturbation of RhoA in a microenvironment-specific manner can be used to promote NP phenotypic expression adding to the knowledge base on the importance of RhoA signaling in NP cells, and the potential to use the RhoA pathway as a therapeutic target. NP cells exhibited a dose response to both CN03 and CT04 treatment, allowing us to screen for an optimal dose for future studies and translation to therapeutics.

The results of this study extend findings about RhoA-specific contractility changes that have been assessed in articular chondrocytes. A study from Hallstrom et. al., shows that RhoA inhibition in 2D increases chondrocyte cell circularity and phenotypic gene expression, mimicking what is observed in our NP cells treated with CT04^23^. CT04-induced inhibition of RhoA in NP cells disrupted F-actin stress-fiber formation and phosphorylation of myosin, blocking actomyosin contractility which led to a rounded cell shape. CT04-treatement in 2D also enriches the ECM related pathways. Accordingly, there was an increase in expression common NP ECM molecule, *ACAN*, as well as *CHST3* which encodes a chondroitin sulfotransferase and has been shown to improve cartilage endplate cell proliferation^33^. *GDF5* was increased, which plays a role in matrix metabolism and NP cell survival^34^. *MUSTN1* was increased which is implicated in chondrogenesis. Conversely, *ADAMTS1* and *ADAMTS9*, both aggrecanases, as well as *COL1* were downregulated in NP cells treated with CT04 in 2D. These data highlight that inhibition of RhoA signaling and retention of rounded cell morphology positively impacts ECM expression and diminished expression of catabolic enzymes. These findings are also consistent with the findings from a prior study by Fearing et. al. that showed retention of rounded NP cell morphology using an F-actin inhibitor (LatB) increases expression of *ACAN* and *COL2*^18^.

NP cells cultured in 2D experience high state of mechanoactivation due to adhesion to tissue culture plastic. Thus, further activation of RhoA using CN03 produced less of an effect than inhibition with CT04 due to the elevated level of baseline contractility. Treatment of chondrocytes in 2D with LPA, a pan-Rho activator, increases F-actin stress fibers and cell area^23^. In our work, CN03 treatment in 2D increased intensity of pMLC and actin, however it did not alter the cellular area. Rho signaling activation has also been linked to chondrocyte dedifferentiation and impaired ECM organization^35,36^. Similarly, NP cells in 2D treated with CN03 exhibit decreased expression of chondrogenesis marker, *GDF5*. With EnrichR analysis, CN03 enriched an ECM organization pathway in part due to the upregulation of two components which have been studied in the ECM: *COL6A6* and *LAMB3. COL6A6* encodes α6(VI) chains which are similar to COL6A3 encoded α3(VI) chains^37,38^. While the specific gene is understudied in the IVD, collagen VI has been found in the pericellular matrix of nucleus pulposus cells^39,40^. Collagen VI levels may elevate with IDD ^41,42^, so the upregulation of COL6A6 could instead be a marker of fibrosis, and therefore differentiation. rather than a pro-NP shift. *LAMB3* encodes for a subunit of [laminin](https://pmc.ncbi.nlm.nih.gov/articles/PMC3332080/)-332 and is enriched in the NP, regulating cell adhesion and spreading via integrin interactions^43^. We also observed increases in expression of *ITGA7*, which is a laminin receptor, with CN03 treatment in 2D, which is consistent with the increased laminin expression^44,45^.

Across both bulk RNAsequencing datasets, we observed changes in the expression of integrin-encoding genes, with opposing patterns depending on RhoA modulation. With RhoA inhibition, *ITGA9* increased and *ITGA8* decreased, whereas RhoA activation had the opposite effect. The increase in *ITGA9* is consistent with prior studies linking it to suppression of RhoA signaling in other systems^46^. Interestingly, *ITGA9* ligands, such as tenascin c (gene: *TNN*) or ADAM15, were also upregulated with CT04 addition^47,48^. In contrast, *ITGA8*, is implicated in contractile cells, such as [smooth](https://www.nature.com/articles/s44161-022-00162-1) muscle cells, which highlights the increase in contraction of the NP cells treated with CN03^49^. It is found that *ITGA8* is a key focal adhesion gene for osteogenesis, potentially highlighting the dedifferentiation of these NP cells in 2D with RhoA activation^50^. Upregulation of *ITGA8* was accompanied by *ITGA7,* which is heavily studied in muscular dystrophy, where overexpression increases the regenerative capacity of muscle^44,45,51^. These results further confirm prior work that highlights that integrins have an influence on the RhoA signaling pathway. To reduce this potential confounding variable, we used alginate hydrogel to culture NP cells in the 3D studies since it is bioinert and lacks cell-binding motifs, thus allowing cell shape and RhoA pathway modulation to be evaluated with reduced influence from focal adhesion engagement at early time points. Alginate is also widely used for NP and chondrocyte culture due to its hydration and tunable mechanical properties to mimic *in situ* microenvironment of these tissues^29,52–54^. While incorporation of RGD motifs to these culture systems may more closely mimic the *in situ* NP cell – ECM interactions, alginate without integrin-binding motifs can be beneficial for chondrogenesis^55,56^.

NP cells in 3D alginate exhibited lower actomyosin contractility than in 2D, observed by the loss of stress fibers and decreased expression of actomyosin genes. This reduction over time enabled evaluation of whether restoring RhoA activity could further promote NP cell roundedness and enhance phenotypic and ECM gene expression. In fact, CN03 treatment at 1-day increases roundedness and *ACAN* gene expression, and at 7-days of daily dosing increases *ACAN* and *KRT19* expression. There are some studies linking the activation of RhoA signaling to sulfated glycosaminoglycan synthesis, through TGFB signaling and the JNK pathway^57^. In the future, long term culture responses to RhoA activation would extend the findings of the current studies using sustained delivery systems.

A key observation is that RhoA activation in 3D and RhoA inhibition in 2D both increase roundedness of NP cells and increase in *ACAN* expression. These data suggest that *ACAN* is sensitive to RhoA modulation and resulting morphological change in NP cells in both 2D and 3D. The link between aggrecan and cell morphology in 2D has been studied in MSCs, where an increase in polygonal cell shape increases aggrecan mRNA expression^58^. As described previously, there is a body of work that highlights that inhibition of contractile elements in chondrocytes or NP cells in 2D promotes aggrecan expression, which we observe in our work with RhoA disruption in 2D^18,23^. However, it remains unclear why activation of RhoA promote expression of *ACAN* in 3D culture. 3D culture alone increases NP cell phenotype, and activation of RhoA further increases cell roundedness based on area and circularity measurements^29,59^. However, RhoA activation and increases in actomyosin contractility are commonly associated with stress fiber formation. In non-adhesive systems lacking focal adhesions, such as in alginate, RhoA activation may promote cell contraction in a different manner than stress fiber assembly, which needs to be further studied. A limitation of this study is that while the change from 2D to 3D culture is a way to change cell morphology, we are also changing the stiffness of the environment, which can also influence RhoA signaling.

This study highlights the microenvironment-specific contexts in which RhoA signaling regulates NP cell morphology, and therefore phenotype. Using CN03 and CT04, we reaffirm the importance of maintaining rounded NP cell morphology to enhance NP cell phenotype and ECM expression. While prior studies from our group have shown that CN03 protects against possible catabolism from NF-kB signaling^4^, the current study shows that CN03 to be a modulator of cell phenotype and ECM production. These findings advance understanding of how microenvironment and RhoA signaling interact to govern cell morphology promote NP phenotype and ECM. Furthermore, Rho/ROCK could be a potential molecular target for the development of IDD therapeutic and regenerative strategies.

## ACKNOWLEDGMENTS

This study was supported in part by National Institute of Arthritis and Musculoskeletal and Skin Diseases (NIAMS) T32AR080744, R01AR077760, R21AR080516. This study used the Confocal and Specialized Microscopy Shared Resource of the Herbert Irving Comprehensive Cancer Center at Columbia University, funded in part through NIH/NCI Cancer Center Support Grant P30CA013696. Research reported in this publication using the ImageStreamX MkII imaging cytometer was performed in the Columbia University Stem Cell Initiative Flow Cytometry core facility at Columbia University Irving Medical Center and was supported by the Office Of The Director, National Institutes Of Health under Award Number S10OD026845. The content is solely the responsibility of the authors and does not necessarily represent the official views of the National Institutes of Health.

