## Supplemental Figure 1 for "Regulation of Nucleus Pulposus Cell Phenotype Through RhoA Signaling and Microenvironment"

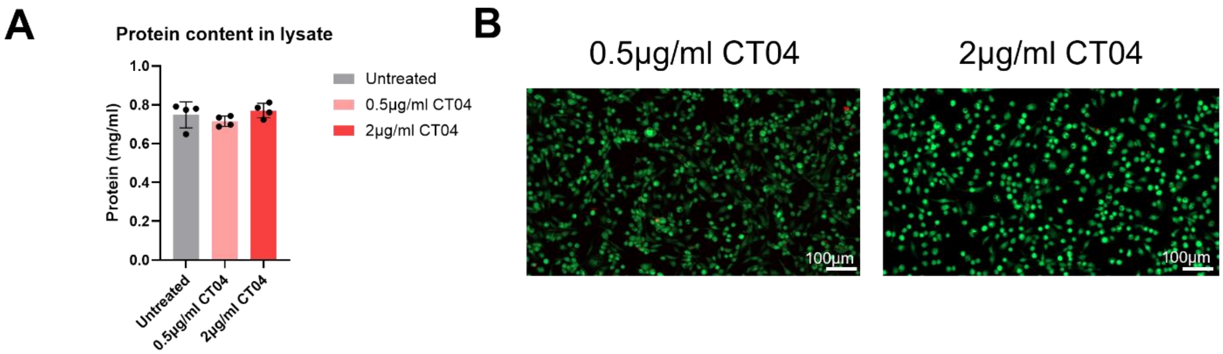


**Figure S1. CT04 treatment does not affect NP cell viability.** A) Protein content in all cell lystates after treatment is consistent. B) Live-dead staining using calcein AM (green) and ethidium homodimer-1 (red) shows that NP cells remain viable after CT04 treatment.
